# Integrated functional genomic analysis identifies the regulatory variants underlying a major QTL for disease resistance in European sea bass

**DOI:** 10.1101/2024.07.06.602346

**Authors:** Robert Mukiibi, Serena Ferraresso, Rafaella Franch, Luca Peruzza, Giulia Dalla Rovere, Massimiliano Babbucci, Daniela Bertotto, Anna Toffan, Francesco Pascoli, Sara Faggion, Carolina Peñaloza, Costas S. Tsigenopoulos, Ross D. Houston, Luca Bargelloni, Diego Robledo

## Abstract

**Background:** Viral nervous necrosis (VNN) is a viral disease threatening the sustainability of global aquaculture, and affecting over 50 of farmed and ecologically important species. A major QTL for resistance to VNN has been previously described in European sea bass, but the underlying causal gene(s) and mutation(s) are unknown. To identify the mechanisms and genetic factors underpinning resistance to VNN, we integrated farmed and wild genetic data with multiple functional genomics assays in a farmed European sea bass population.

**Results:** A high heritability (h2 ∼ 0.40) was estimated for VNN resistance. A major QTL for this trait was confirmed on chromosome 3, and whole-genome resequencing narrowed its location to a small region containing 4 copies of interferon alpha inducible protein 27-like 2A (*IFI27L2A*) genes, and one copy of the interferon alpha inducible protein 27-like 2 (*IFI27L2*) gene. RNA sequencing revealed a clear association between the QTL genotype and the expression of two of the *IFI27L2A* genes, and the *IFI27L2* gene. Integration with chromatin accessibility and histone modification data pinpointed two SNPs in active regulatory regions of two of these genes (*IFI27L2A* and *IFI27L2*), and transcription factor binding site gains for the resistant alleles were predicted. These alleles, particularly the SNP variant CHR3:10077301, exhibited higher frequency in Eastern Mediterranean sea bass populations, which show considerably higher levels of resistance to VNN.

**Conclusions:** The SNP variant CHR3:10077301, through modulation of *IFI27L2* and *IFI27L2A* genes, is likely the causative mutation underlying resistance to VNN in European sea bass. This is one of the first causative mutations discovered for disease resistance traits, and paves the way for marker-assisted selection as well as biotechnological approaches to enhance resistance to VNN in European sea bass and other susceptible species.

## BACKGROUND

The European sea bass (*Dicentrarchus labrax*) is a highly esteemed marine fish in Europe and the Mediterranean, boasting major economic and cultural value(Vandeputte *et al*. 2019). Over the past two decades, global aquaculture production of European sea bass has experienced remarkable growth, steadily increasing from 7,694 tons in 2000 to 299,810 tons in 2021; the major producing countries are Turkey, contributing approximately 52%, and Greece, accounting for around 17% of the total global production(FAO 2024). Aquaculture dominates European sea bass production, constituting 98% of the total, as wild fisheries production has witnessed a steady decline of 38% over the past decade(FAO 2024). However, the industry faces an increasing challenge in the form of infectious diseases, which pose a significant threat to both the sustainability of the industry and the welfare of farmed European sea bass. Infectious diseases currently account for approximately 10% of all fish mortalities on aquaculture farms(Muniesa *et al*. 2020).

Viral nervous necrosis (VNN) stands as the main viral infectious disease, currently responsible for 15% of total on-farm infectious disease-related mortalities in European sea bass(Muniesa *et al*. 2020). Beyond the direct toll of mortalities, sea bass producers also suffer economic losses stemming from fish that survive the viral infection but experience delayed growth(Barsøe *et al*. 2021). Primarily affecting larvae and juvenile fish, VNN inflicts moderate to high mortality rates, reaching up to 100% in affected farms(Le Breton *et al*. 1997). The causative agent of VNN, Nervous Necrosis Virus (NNV), belongs to the *Betanodavirus* genus in the *Nodaviridae* family, and is a non-enveloped single positive-stranded RNA virus(Chi *et al*. 2016). The main target of NNV is the central nervous system, particularly the brain, spinal cord and the retina, where viral replication and multiplication occurs(Chi *et al*. 2016). NNV has a broad host range, as it can infect a wide range of aquaculture fish species such as Asian seabass (*Lates calcarifer*), greasy grouper (*Epinephelus tauvina*), red-spotted grouper (*Epinephelus akaara*), gilthead sea bream (*Sparus aurata*), and turbot (*Scophthalmus maximus*)(Chi *et al*. 2016; Vandeputte *et al*. 2017; Yang *et al*. 2022). Effective prevention and treatment strategies for VNN remain elusive, and therefore current practices focus on minimizing disease outbreaks through stringent biosecurity measures, vaccination, and developing experimental therapeutics(Chi *et al*. 2016; Yang *et al*. 2022).

Recent studies have revealed significant phenotypic and genetic variation in resistance to VNN within farmed European sea bass populations. Moderate to high heritability estimates of 0.10 to 0.43 have been reported for this trait(Vandeputte *et al*. 2017; Palaiokostas *et al*. 2018; Faggion *et al*. 2021; Griot *et al*. 2021; Vela-Avitúa *et al*. 2022), indicating the potential for selective breeding to develop strains with heightened genetic resistance to the virus. Notably, a major quantitative trait locus (QTL) with a large impact on VNN resistance has been identified on chromosome 3 (previously LG22) of the sea bass genome in three distinct farmed populations(Palaiokostas *et al*. 2018; Griot *et al*. 2021; Vela-Avitúa *et al*. 2022; Delpuech *et al*. 2023). This major QTL accounts for over 33% of the genetic variation in resistance to VNN(Palaiokostas *et al*. 2018; Griot *et al*. 2021; Vela-Avitúa *et al*. 2022; Delpuech *et al*. 2023), thus marker-assisted selection can drive rapid and significant improvement in this trait, leading to sustained reductions in disease outbreaks.

Moreover, understanding the genomic basis underlying this QTL would open avenues for leveraging genome engineering approaches to bolster VNN resistance in populations and species lacking genetic variation at this resistance locus. Several genes, including *PLK4, HSPA4L and REEP1,* have been suggested as potential genes underlying VNN resistance due to their proximity to the QTL and their recognised antiviral function(Vela-Avitúa *et al*. 2022). In a recent large study by Delpuech *et al*. (2023) on a large population, the putative causative mutation was proposed to reside approximately 1.9 and 6 kb downstream of the *ZDHHC14* and IFI27-like genes on chromosome 3, the same chromosome/linkage group where previous QTLs for the same trait were reported, albeit at a considerable distance(Delpuech *et al*. 2023). In the same study, other less prominent QTLs were identified on the same chromosome 3. It is worth noting that all these studies were conducted on a highly fragmented genome assembly(Tine *et al*. 2014), and so far no functional evidence in support of any causative gene has been reported.

The availability of highly contiguous and well-annotated genomes is essential to uncover the causative variants underlying QTL(Clark *et al*. 2020). In recent years, new high-quality genome assemblies have been released for many aquaculture species(Martínez *et al*. 2021; Etherington *et al*. 2022; de la Herrán *et al*. 2023), including the European sea bass (GCA_905237075.1, contig N50 = 12 Mb). However, the functional annotation of these assemblies is mostly limited to gene positions along the genome, facilitated by the abundant collection of RNA sequencing datasets for many teleost species and the general conservation of coding sequences. Considering the key role of regulatory variants in shaping the genetic architecture of complex traits (Emilsson *et al*. 2008; Albert & Kruglyak 2015), it is crucial to extend the functional annotation to the non-coding regions of the genome to advance toward the identification of causative variants underlying QTLs.

Recent studies, such as large genotype-tissue expression analyses in human(Consortium 2020), cattle(Liu *et al*. 2022) and pig(Teng *et al*. 2024), have identified expression QTLs (eQTLs) across different tissues. Notably, some of these eQTLs co-localize with QTL regions strongly associated with complex traits of both health and economic or production significance. This evidence underscores the importance of identifying loci modulating gene expression (eQTLs) and colocalizing with disease resistance QTLs, which can significantly contribute to unraveling the genetic architecture of complex traits, including disease resistance to pathogenic infections. Additionally, novel tools to identify and profile the regulatory regions of the genomes have become available in recent years, such as the assay for transposase-accessible chromatin sequencing (ATAC-seq) and the chromatin immunoprecipitation sequencing (ChIP-seq). Integrating multiple –omic evidence with QTL/eQTL data is pivotal for the reliable identification of causal genetic variants underlying variability in a complex trait, however these datasets are scarce non-model species.

In this study, we investigated the genetic architecture and underlying functional genomic basis of resistance to VNN in a non-model species, the European sea bass. By combining whole-genome genotyping, targeted eQTL identification, analysis of chromatin accessibility and ChIP-seq data, and population genomic evidence, we unveiled reliably the putative mechanisms driving a major QTL for resistance to VNN in this species. This integrative approach provides a comprehensive understanding of the genetic factors contributing to disease resistance and sheds light on potential avenues for future research and breeding strategies in non-model organisms like the European sea bass.

## RESULTS

### VNN infection challenge, survival analysis and population genetic structure

No fish mortality was observed during the first 5 days of the VNN challenge experiment; however, the peak of mortality was observed on day 6, and then the number of deaths decreased steadily reaching zero on day 29, when the experiment was terminated (Figure 1A). The overall mortality of the challenge experiment was 46.84% (Figure 1A), close to that observed in previous studies with similar experimental design(Faggion *et al*. 2021; Faggion *et al*. 2022).

**Figure 1.**
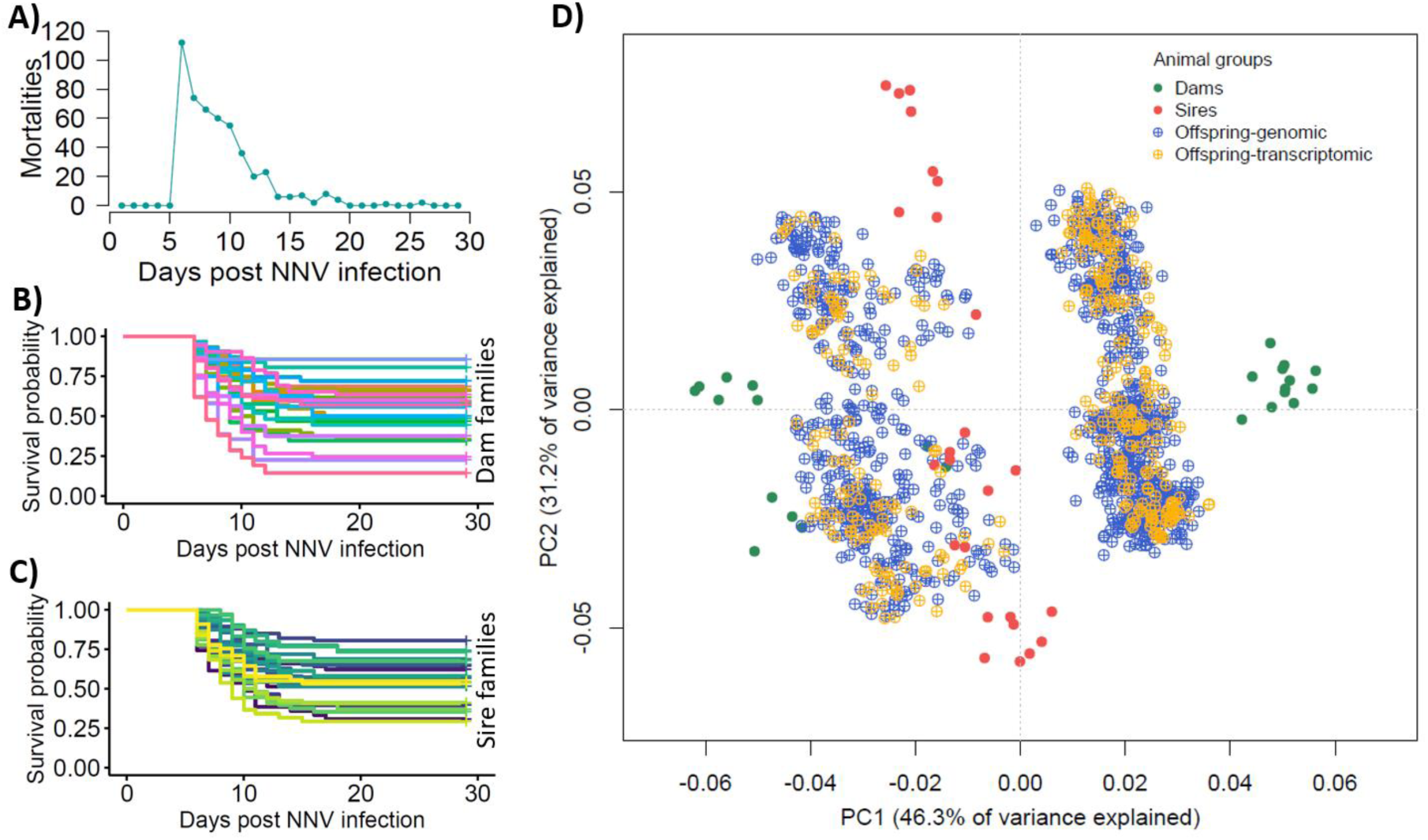
VNN challenge experiment results and genetic structure of the European sea bass challenged population. A) Number of mortalities each day of the challenge experiment; B) Survival curves for each dam family throughtout the experiment; C) Survival curves for each sire family throughout the experiment; D) Principal component analyses showing the genetic structure of the study population.

All challenged animals and their parents were genotyped using a 30K SNP array, and then imputed to whole-genome genotypes from 90 whole-genome sequenced individuals (all 50 parents and 40 offspring). Genotypes were used to reconstruct the pedigree. Each sire and dam contributed a substantial number of offspring to the challenge population (Supplementary file 1), and a survival analysis showed phenotypic variation in resistance to VNN infection between both dam and sire families, with overall survival ranges of 14.3 – 85.7% for dam families (Figure 1B), and 29.2 – 80.6% for sire families (Figure 1C).

Principal component analysis of parents and offspring showed important genetic structure, likely driven by the geographic origin of the parents and specially of dams (Figure 1D). In fact, the base population for the breeding programme at the hatchery contributing the biological material for the present study was established mixing European sea bass of Atlantic and Mediterranean origin populations, which have been consistently reported to show significant genetic divergence (Tine *et al*. 2014).

### Genetic parameters and association study

Heritability estimates for VNN resistance were moderate to high, ranging between 0.35 and 0.45 (Table 1). These estimates are in general higher than those reported in previous studies using different farmed sea bass populations(Palaiokostas *et al*. 2018; Faggion *et al*. 2021; Griot *et al*. 2021; Vela-Avitúa *et al*. 2022; Delpuech *et al*. 2023). Heritabilities were slightly higher when calculated using whole-genome genotypes, and the heritability of days to death (DD) was higher than for binary survival (BS) (Table 1). There was high genetic correlation between BS and DD (0.99, Table 1), therefore both measurements represent exactly the same trait.

**Table 1.** Heritabilities and genetic correlation for resistance to VNN; days to death (DD) and binary survival (BS).

| | $h^2 \pm se$ (BS) | $h^2 \pm se$ (DD) | $r_g \pm se$ (BS & DD) |
| --- | --- | --- | --- |
| **SNP array** | $0.35 \pm 0.05$ | $0.39 \pm 0.05$ | $0.99 \pm 0.00$ |
| **Imputed WGS** | $0.40 \pm 0.06$ | $0.45 \pm 0.06$ | - |

GWAS analysis for resistance to VNN revealed a major QTL in chromosome 3 (Figure 2A). The significant markers explained up to 15.27% of the phenotypic variance, and up to 38.28% of the genetic variance in survival to VNN (Figure 2B, Supplementary File 2). The use of whole-genome sequencing improved the power of these analyses, as the variants explaining the largest percentage of genetic variance were not present in the SNP array(Mukiibi *et al*. 2022). Consistently with the high percentage of variance explained, the QTL genotype had a large and additive impact on survival rate, as fish homozygous for the resistance genotype showed 88.2 - 89.8% survival, heterozygous individuals 59.6 – 61.0%, and homozygous animals for the susceptible genotype 32.2 – 32.6% (Figure 2C). Homozygous resistant fish showed ∼55% increased survival compared to homozygous susceptible fish.

**Figure 2.**
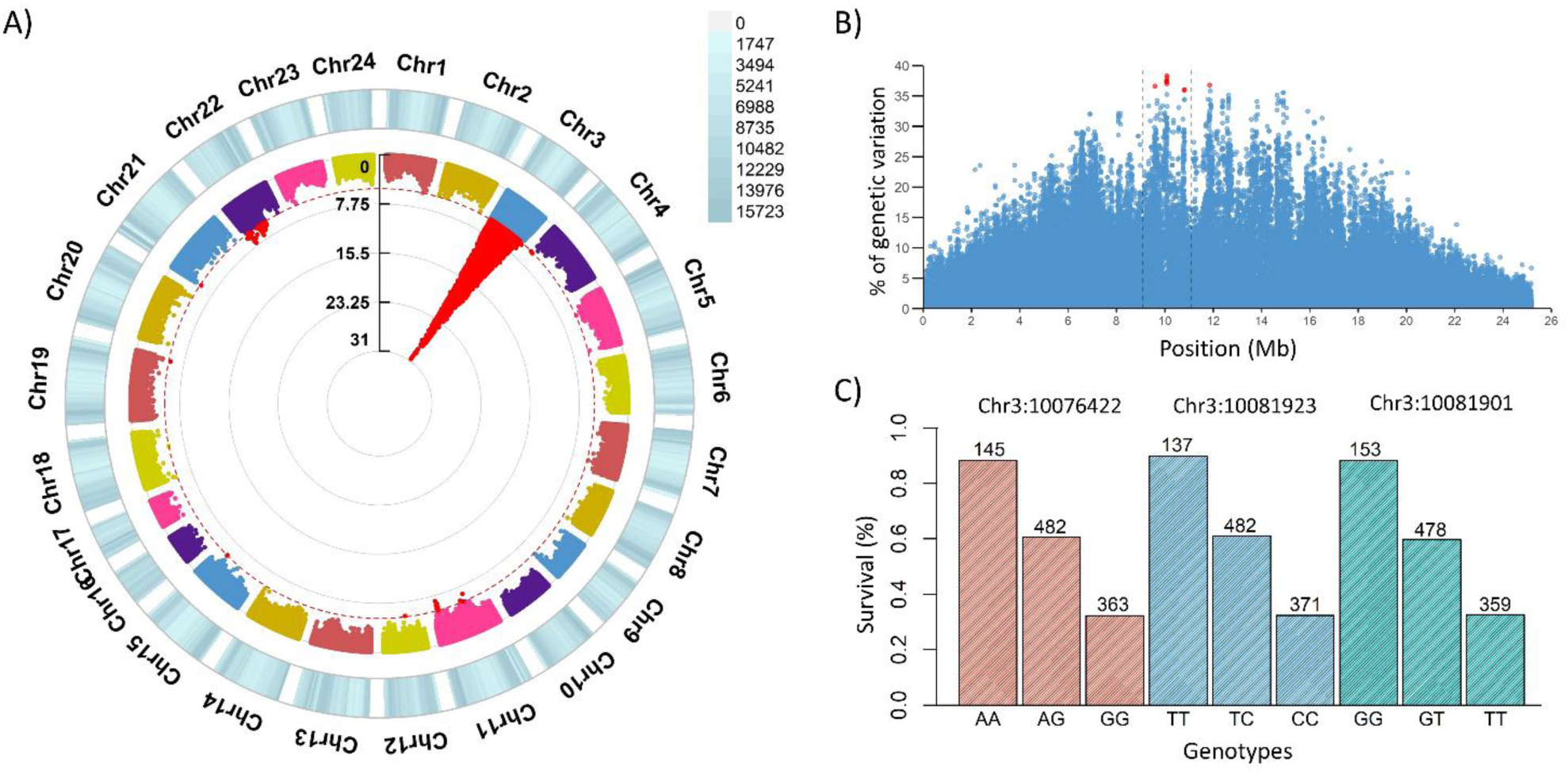
Signatures of resistance to VNN in the European sea bass genome. A) Circular Manhattan plot with the results of the GWAS for VNN resistance, showing SNP density along each chromosome; B) Percentage of genetic variance explained by the SNPs on Chromosome 3; C) Barplot showing the percentage of survival of the animals with each genotype for the three most significant SNPs.

### Annotation of the QTL region

While the SNPs explaining >34% of the genetic variance span a region >1.5 Mb, the most significant variants are located in a small region of approximately 10 kb. This region is characterised by the presence of multiple genes annotated as interferon alpha inducible protein 27-like 2A (*IFI27L2A*), and an interferon alpha inducible protein 27-like 2 (*IFI27L2*) gene (Figure 3). However, the two available genome annotations (dlabrax2021.112 from Ensembl and GCF_905237075.1-RS_2023_02 from NCBI) for the current European sea bass genome assembly (GCA_905237075.1) show slight differences in this QTL region on chromosome 3. The 5’ untranslated region (UTR) of ENSDLAG00005028932 (Ensembl) is extremely long, while this 5’UTR sequence is replaced by a novel gene (LOC127358630) in the NCBI annotation, which is not present in the Ensembl annotation. On the contrary, a gene model present in the Ensembl annotation (ENSDLAG00005026832) is missing in the annotation of the same region by NCBI. Full-length sequence RNA-seq generated for sea bass head kidney and brain were used to establish the correct annotation for the region (Figure 3), suggesting the presence of both LOC127358630 and ENSDLAG00005026832 genes. Therefore, a modified version of the NCBI annotation, with the ENSDLAG00005026832 added, was used for all subsequent analyses. In this annotation, there are five different interferon-induced genes (four *IFI27L2A*, and one *IFI27L2*) in the region, all located consecutively in the genome. Upstream of these genes there is a locus coding for galectin-8 (LGALS8, LOC127358615), and downstream a locus encoding a Cytosolic 5’-Nucleotidase 1A (cN-1A, LOC127358605). The variants showing the highest association with resistance to VNN overlap the three most upstream of these interferon-induced genes (LOC127358630, LOC127358685, and ENSDLAG00005026832) (Figure 3).

**Figure 3.**
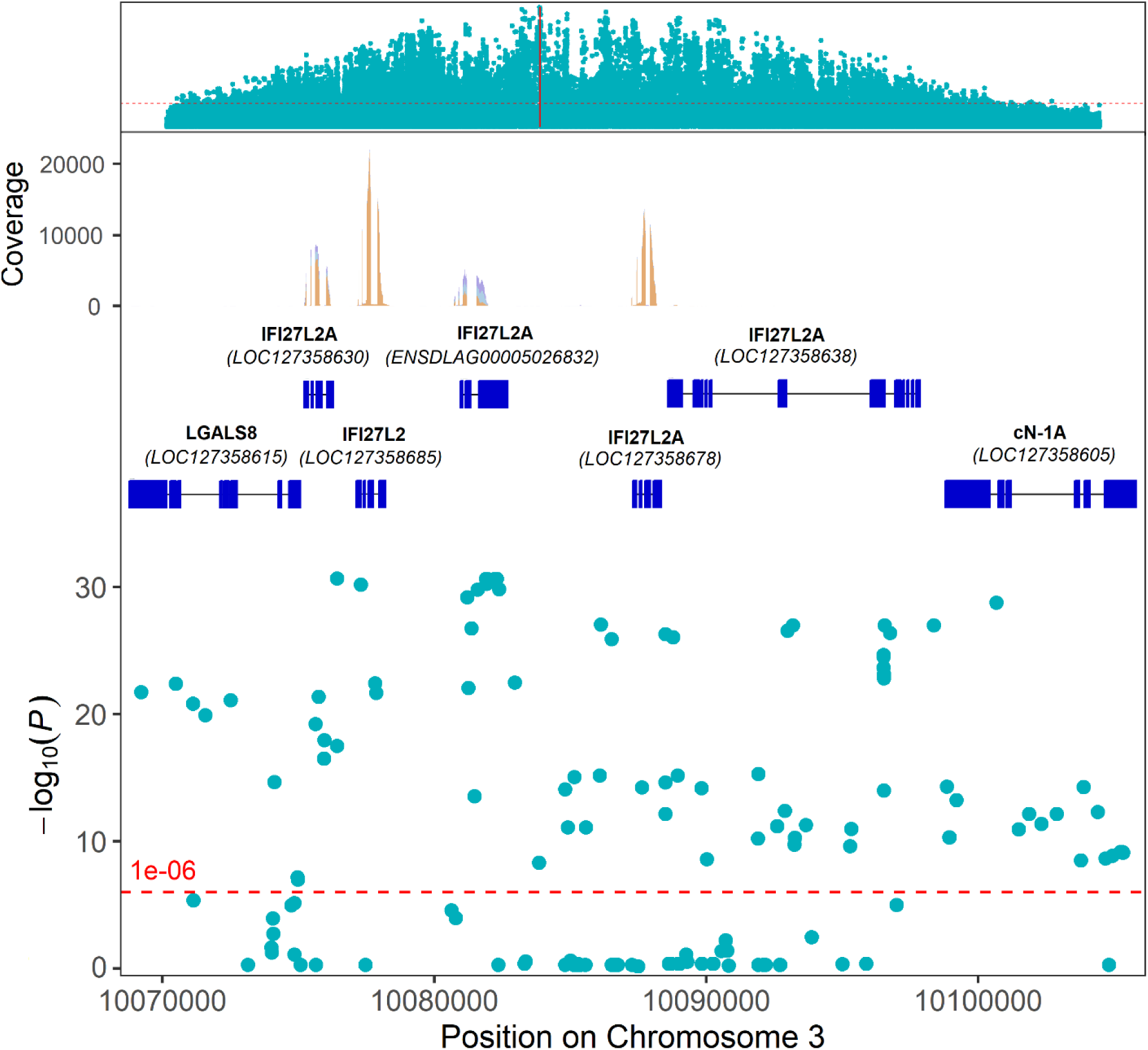
Functional annotation of the VNN resistance major QTL region in European sea bass chromosome 3. The region containing the SNPs showing the highest association with VNN resistance (Chromosome 3: 10068810-10105810) is shown. The figure shows the significance of the SNPs for resistance to VNN across the whole chromosome 3 (Track 1) and across the narrow QTL region (Track 4), as well as the genes present in the region (NCBI annotation + ENSDLAG00005026832, Track 2) and expression evidence of these genes according to head kidney and brain RNA-sequencing (Track 3).

### Evolutionary conservation of the QTL region

To discard potential misassemblies and confirm the existence of multiple copies of interferon-induced genes (*IFI27L2A* and *IFI27L2*), the European sea bass major QTL region was compared with homologous genomic regions in the striped sea bass (*Morone saxatilis)*, a closely related species of the same family Moronidae, and the gilthead sea bream (*Sparus aurata)*, a more distantly related species. Both species have a cluster of four copies of interferon-induced genes (three *IFI27L2A* and one *IFI27L2*), and similar to European sea bass, this gene cluster is flanked by galectin-8 upstream and cytosolic 5’-nucleotidase 1A downstream. Comparison of the homologous genomic regions in the striped bass and the European sea bass showed high sequence conservation and consistent gene order, apart from the absence of a homologue for LOC127358638 in *M. saxatilis* (Figure 4A).

**Figure 4.**
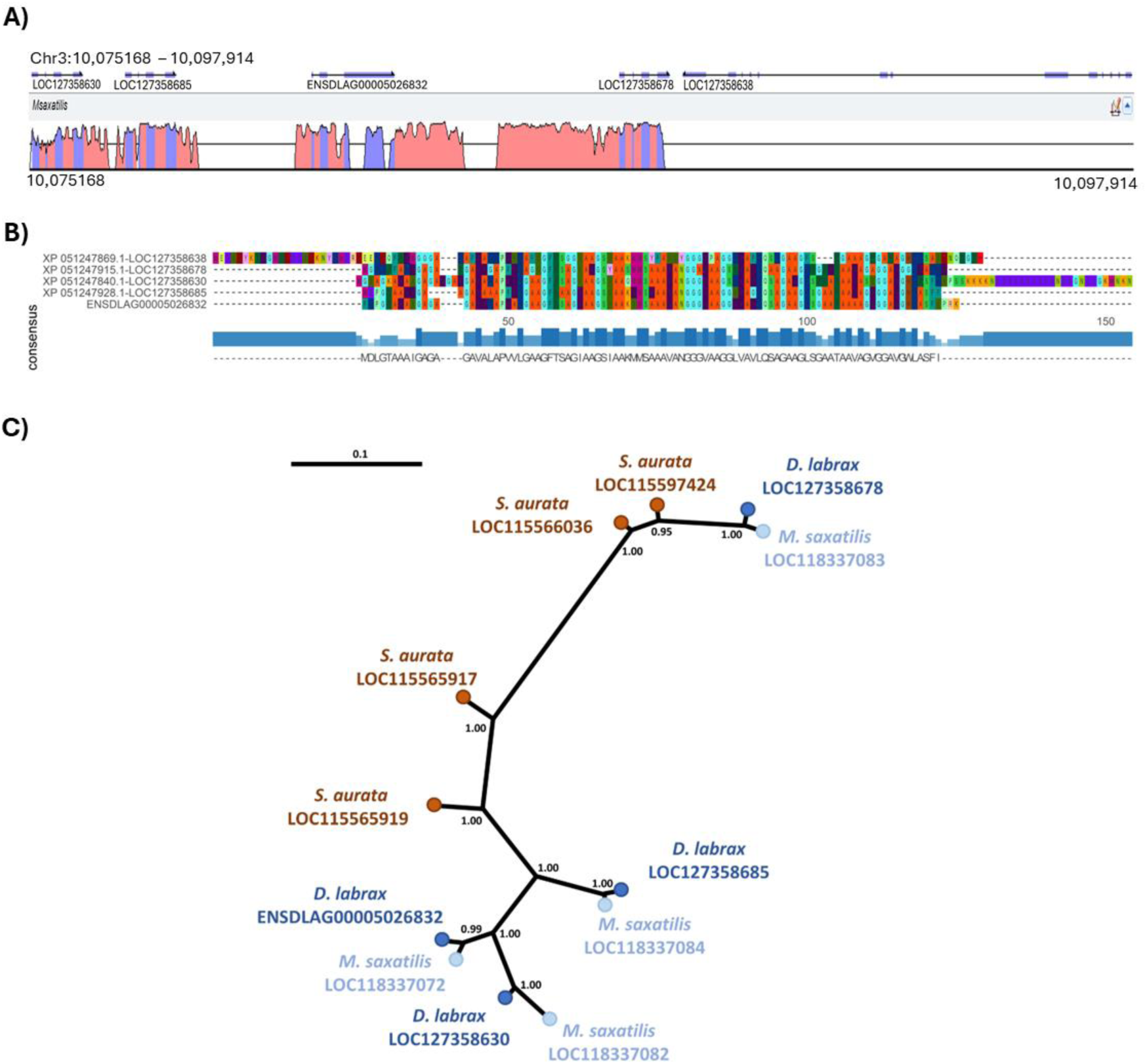
Evolutionary conservation of the major QTL region for VNN. A) “Peaks and Valleys” graph showing the percentage of conservation of the European sea bass VNN resistance QTL region (Chromosome 3: 10,075168 – 10,097,914) with the homologous region in *M. saxatilis* (NCBI NW_023339873.1: 15,596,712 – 15,606,764), “peaks” indicate high similarity between the two species, while “valleys” indicate low similarity; B) Multiple sequence alignment plot of the Amino acid sequences of the five interferon-induced genes in the VNN resistance QTL region of European sea bass, where letters correspond to the standard abbreviation of amino acid residues in the peptide sequences, while “-“ represents a gap in the alignments; C) IFI27L2-like gene tree reconstructed using Bayesian inference. GeneIDs are reported at the branch tips for consistency throughout the document, but the corresponding coding mRNA sequences were used for the analysis (*D. labrax*: LOC127358678 - XM_051391955.1, LOC127358685 - XM_051391968.1, LOC127358630 - XM_051391880.1; *S. aurata*: LOC115566036 - XM_030391784.1, LOC115597424 - XM_030443305.1, LOC115565917 - XM_030391575.1, and LOC115565919 - XM_030391576.1; *M. saxatilis*: LOC118337083 - XM_035673933.1, LOC118337072 - XM_035673917.1, LOC118337084 - XM_035673934.1, LOC118337082 - XM_035673932.1). Numbers at nodes are posterior probability values.

Sequence alignment of the five interferon-induced European sea bass genes/proteins (four *IFI27L2A*, and one *IFI27L2*) revealed that this LOC127358638 has diverged significantly of the other four interferon-induced genes in the cluster, which had relatively high sequence similarity (Figure 4B). Considering the high sequence divergence, and the lack of an homologous gene in *M. saxatilis*, this gene was excluded from phylogenetic analyses since it would significantly disrupt protein alignments. While not as extreme, the other four sea bass interferon-induced genes also showed substantial sequence divergence, with LOC127358678 being quite different from the three upstream interferon-induced genes (Figure 4B).

To reconstruct the evolution of the other four IFI27L2 genes in *D. labrax*, their coding mRNA sequences were aligned against those of *M. saxatilis* and *S. aurata*, which confirmed that these European sea bass interferon-induced genes (LOC127358630, LOC127358685, ENSDLAG00005026832, LOC127358678) have one-to-one orthologues in the striped sea bass genome (Figure 4C), suggesting that duplication events occurred in the common ancestor of the two species. These three genes formed a relatively tight phylogenetic cluster, reflecting their high sequence similarity.

### Expression QTL analysis shows evidence of genetic variants controlling the expression of the three upstream IFI27L2 genes

Brain and head kidney RNA-Seq data of 110 mock and 214 VNN-challenged fish from the same families used for the larger GWAS disease challenge were produced. Of these fish, 322 had acceptable high-quality ∼30K SNP array genotypes and hence imputed to whole-genome genotypes based on the whole-genome resequencing of their parents and 40 sibs. Animals were classified based on their QTL genotypes, and differential expression analyses between individuals homozygous for the resistant allele and homozygous for the susceptible allele (based on the three most significant SNPs) revealed differences for three interferon-induced genes (LOC127358630, LOC127358685, and ENSDLAG00005026832) (Table 2). However, these differences were unstable (i.e., none of the three genes showed significant differences across the two conditions and the two tissues). Nonetheless, the homozygous “resistant” group showed higher expression at all three loci in at least one tissue/condition with statistically significant fold changes (FC) ranging from 1.58 to 2.57 (Table 2).

**Table 2.**
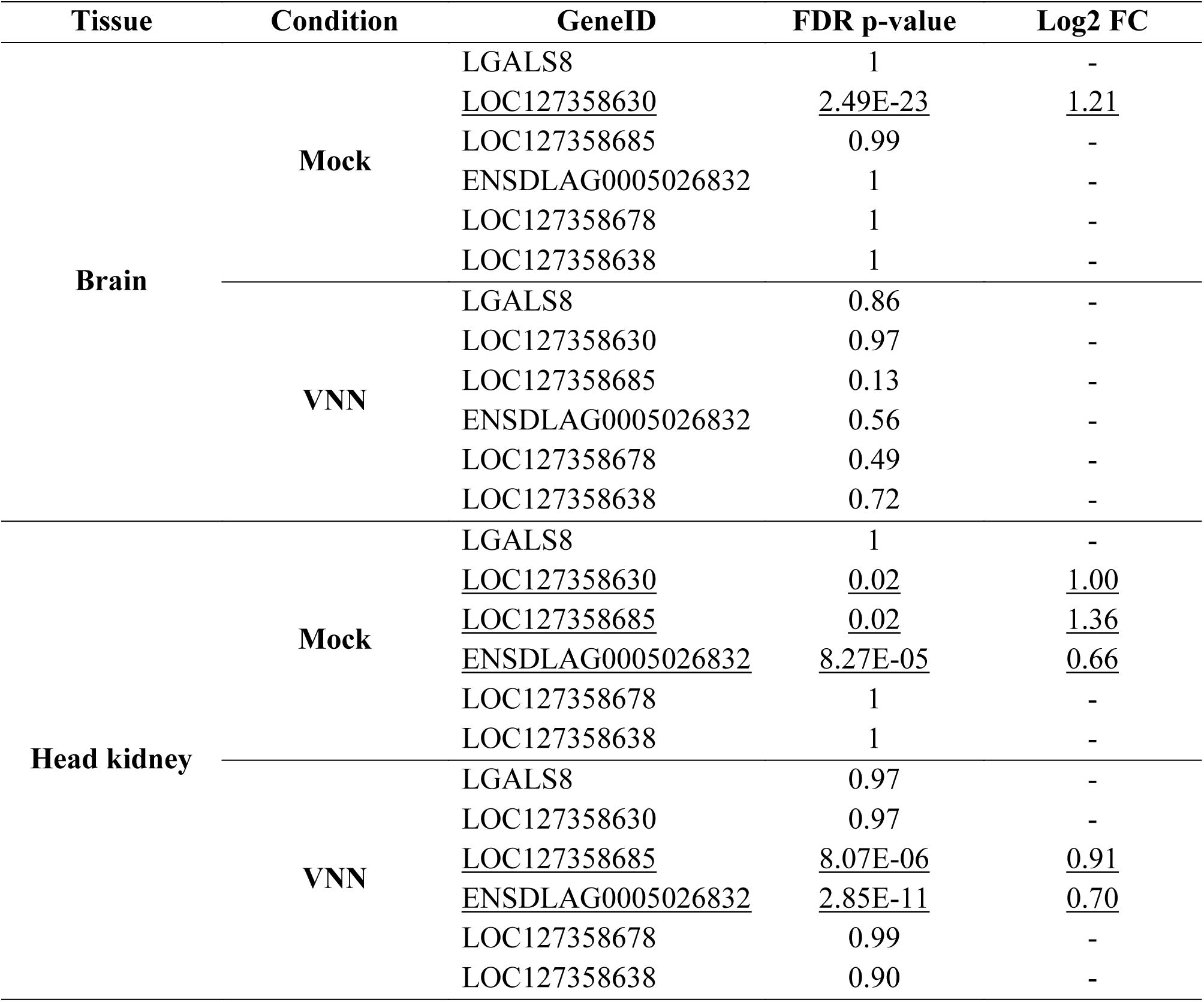
Differential expression between homozygous resistant (RR) and homozygous susceptible (SS) European sea bass fish based on the major VNN resistance QTL genotype.

Consistently, global correlation analyses showed strong correlations (*r* = 0.2 – 0.49; p < 0.005) between the expression of the three interferon-induced genes and predicted VNN resistance breeding values. Noteworthy, the correlations of the three genes varied between conditions and tissues with only LOC127358685 and ENSDLAG0005026832 showing consistent significant correlations in the head kidney in the mock and VNN-challenged fish. Expression QTL (eQTL) analyses for the three interferon-induced genes revealed a clear association between the SNPs in the VNN QTL region and the expression of these genes (Figure 5A). As in the differential expression analyses and correlation analyses, the eQTL results were varied across conditions and tissues, nevertheless, the association between QTL genotypes and the expression of the three interferon-induced genes was unequivocal. Notably, the genetic variants that were highly significant in the eQTL analysis overlapped with those described above associated with VNN resistance(Figure 5B). In all cases, the minor frequency “alternative” allele was associated with higher gene expression as well as better survival after VNN infection, and it showed a clear additive effect, with heterozygous animals showing intermediate expression of IFI27L2 (Figure 5C).

**Figure 5.**
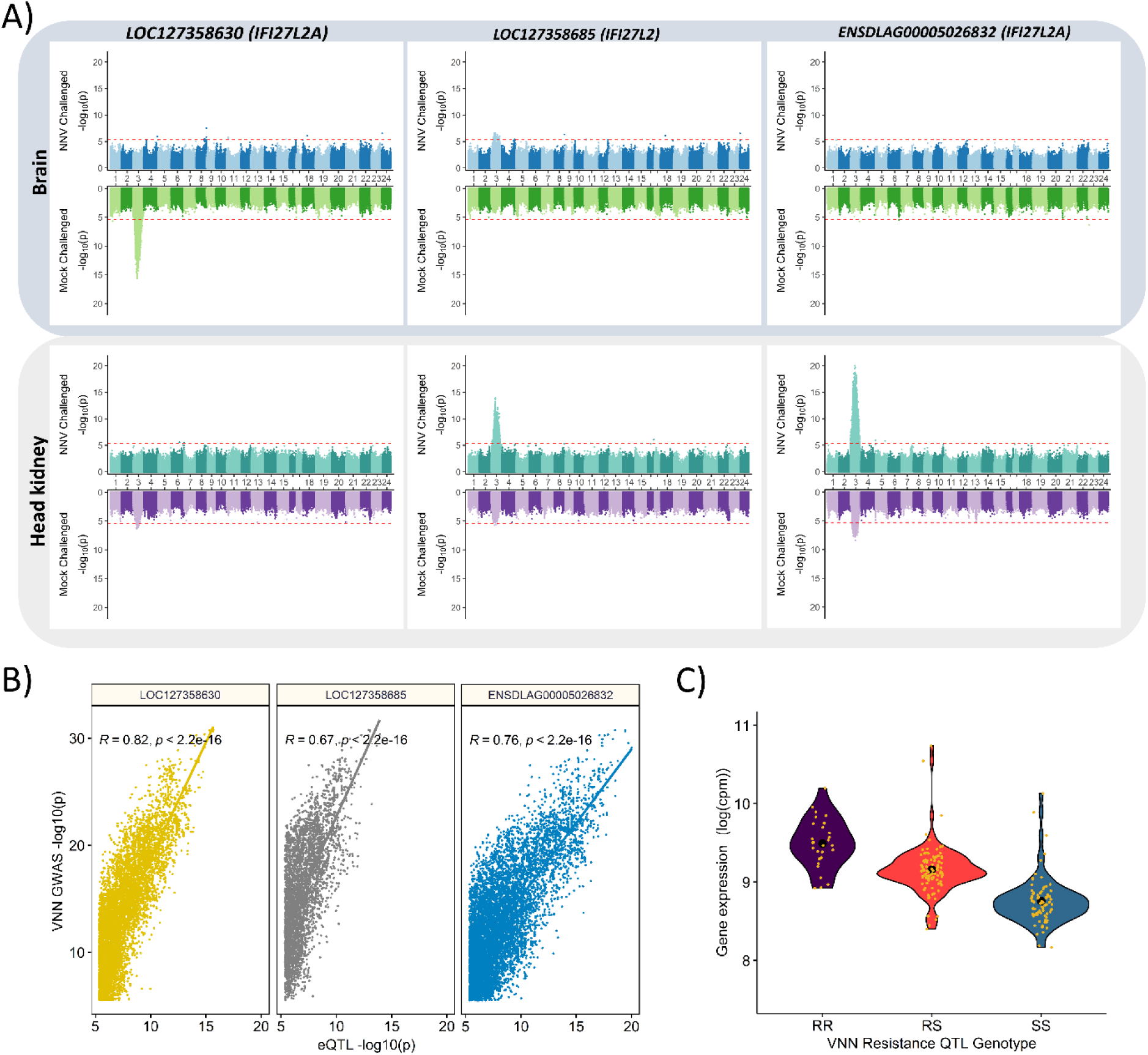
Association between IFI27L2 gene expression in European sea bass and the QTL genotype. A) Expression QTL analysis for the three IFI27L2/2A genes in the proximal region of the VNN resistance QTL in chromosome 3; B) Pearson correlation between the expression QTL analysis p-values for the three proximal IFI27L2/2A genes and the p-values for resistance to VNN based on the whole-genome GWAS, C) violin plot showing the gene expression of IFI27L2/2A in animals with different VNN resistance QTL genotypes: RR (two resistant alleles), RS (one resistant and one susceptible allele), SS (two susceptible alleles).

### **A**nalysis of sequence variants in putative regulatory regions

A subset of 20 fish (10 mock- and 10 VNN-infected) of the ones with RNA-seq data were used to assess chromatin accessibility using ATAC-seq in brain and head kidney tissues. In the major QTL region, four open chromatin regions were identified in the head kidney and three in the brain, with substantial overlap between the tissues (Figure 6). No differences were observed between mock and VNN-challenged samples. Additionally, active regulatory regions were predicted using both ATAC-seq and ChIP-seq for three different histone marks, finding overlap between the open chromatin regions and active regulatory elements (TSS flank and TSS active) (Figure 6). Two single nucleotide variants, located in CHR3:10077301 and CHR3:10081209, showing high association with survival to VNN and with the expression of the three upstream *IFI27L2* genes, were located within the putative regulatory regions of LOC127358685 and ENSDLAG00005026832, respectively, which are shared between brain and head kidney (Figure 6).

**Figure 6.**
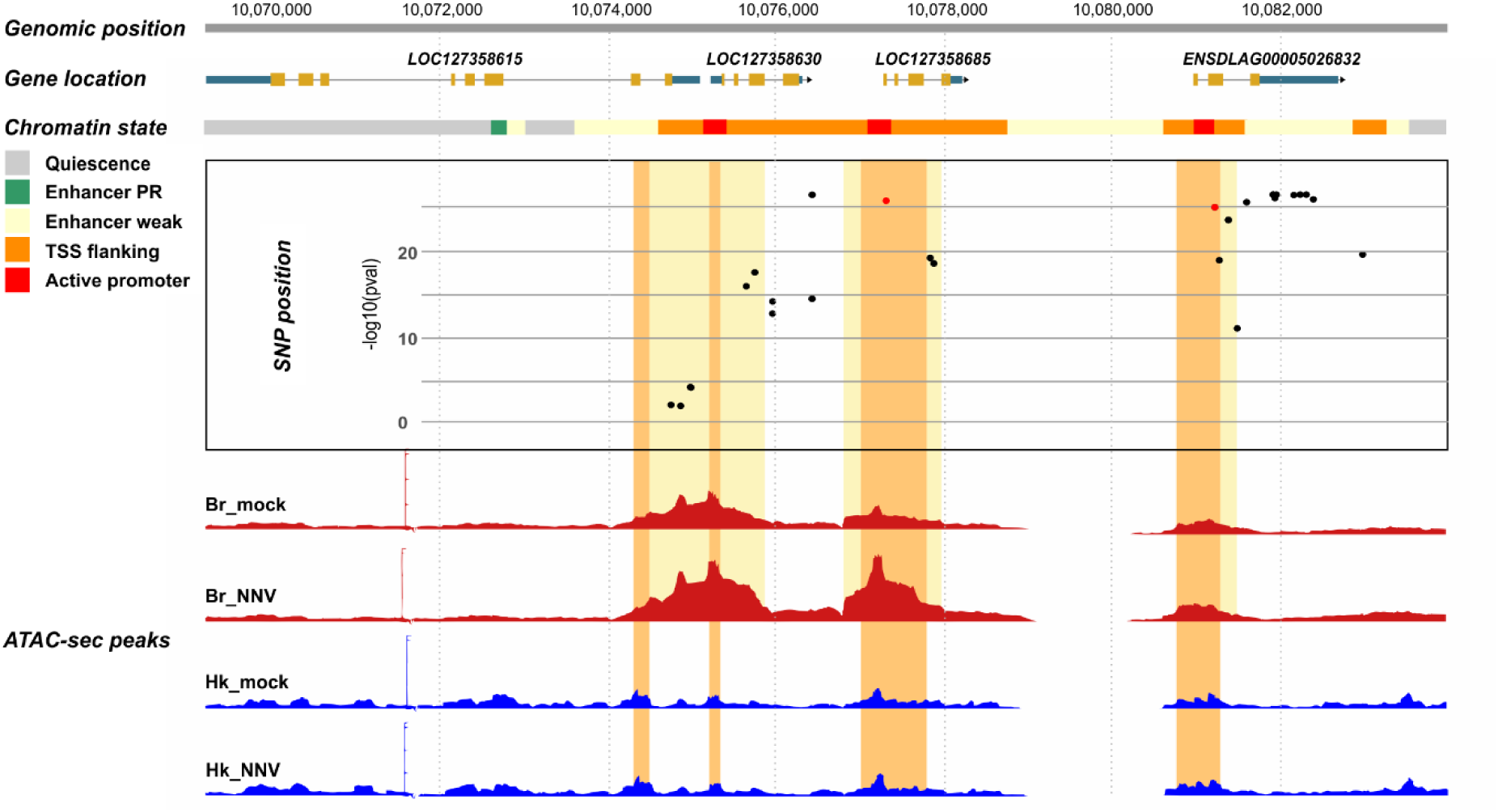
Regulatory landscape of the VNN resistance QTL region in the head kidney and the brain. The figure shows the VNN resistance QTL region, the position of the four expressed IFI27L2 genes, the chromatin state based on ATAC-seq and ChIP-seq assays, VNN resistance GWAS p-values for the SNPs in the regions, and chromatin accessibility peaks in mock and VNN challenges brain (Br) and head kidney (Hk).

To evaluate the potential effect of these two variants on transcriptional regulation, the variant effects on transcription factor binding significant SNPs were predicted using FABIAN-variant. For SNP CHR3:10077301, the alternative (resistant) allele was predicted to lead to a transcription factor binding site (TFBS) gain for human KLF transcription factor 15 (KLF15, ENSG00000163884) with a high score (0.8146 on a 0-1 scale). For CHR3:10081209, two TFBS gains were predicted for the alternative (resistant) allele, for zinc finger protein 140 (ZNF140, ENSG00000196387) and TEA domain transcription factor 4 (TEAD4, ENSG00000197905), respectively, with a score of 0.9808 and 0.9421. Human ZNF140 does not have an orthologue in the European sea bass, while there are 1:1 orthologues for KLF15 (ENSDLAG00005009091) and TEAD4 (ENSDLAG00005022250). Sequence similarity at the protein level was high for TEAD4 (on average >80%) and very high for the TEA domain. For KLF15, similarity was moderate (on average >50%), although the DNA binding domain was highly conserved.

### Distribution of allele frequencies across natural populations

The two variants were also analysed in terms of allele frequencies in wild populations of sea bass from different geographical areas (i.e., Atlantic Ocean, Western Mediterranean Sea, Eastern Mediterranean Sea), which are known to be genetically divergent and show differential resistance to VNN; Eastern Mediterranean populations have a significantly higher survival (up to three times) upon experimental infection with VNN compared to fish of Atlantic and Western Mediterranean origin [4]. Analysis of published pool-sequencing WGS data (Peñaloza et al. 2021) allowed the estimation of allele frequencies for the two variants (Table 3). The minor allele variants associated with disease resistance and higher expression of interferon-induced genes showed significantly higher frequency in Eastern Mediterranean population samples (CHR3:10077301 p<0.00001; CHR:10081209 p<0.01), with the difference substantially larger for CHR3:10077301 (0.45 vs 0.08).

**Table 3.** Minor allele frequencies for the two main VNN resistance candidate SNPs.

| *Variant* | *ATL* | *WMED1* | *WMED2* | *EMED1* | *EMED2* |
| --- | --- | --- | --- | --- | --- |
| CHR3:10077301 | 0.21 | 0.15 | 0.25 | 0.55 | 0.77 |
| CHR3:10081209 | 0.05 | 0.07 | 0 | 0.12 | 0.13 |

The minor alleles are the VNN resistance alleles. ATL – Atlantic population (susceptible), WMED1 & WMED2 – Western Mediterranean populations (susceptible); EMED1 & EMED2 (resistant).

## DISCUSSION

Our results confirm that VNN resistance has moderate to high heritability (0.35 to 0.45) in farmed European sea bass, with slightly higher values than suggested by previous studies(Palaiokostas *et al*. 2018; Griot *et al*. 2021; Vela-Avitúa *et al*. 2022), and therefore there is great potential for selective breeding to produce fish that are more resistant to this disease, reducing VNN’s economic impact and improving fish welfare. Our GWAS results also confirmed the presence of a very strong QTL for VNN resistance, located on Chromosome 3 of the European sea bass genome, consistent with similar studies in different farmed sea bass populations(Palaiokostas *et al*. 2018; Griot *et al*. 2021; Vela-Avitúa *et al*. 2022). The most significant SNPs in the QTL explained up to 38% of the total genetic variance, which is comparable to the report by Vela-Avitúa *et al*. (2022) where the most significant SNP explained 33.33% of VNN resistance genetic variance(Vela-Avitúa *et al*. 2022). However, Griot et *al.* (2021) reported a substantially lower genetic variance (9%) explained by the most significant markers(Griot *et al*. 2021). This difference could depend on the infection challenge model; while in both our study and Vela-Avitúa *et al*. (2022), fish were infected via intraperitoneal/intramuscular injection. Griot et *al.* (2021) performed an immersion challenge, which could result into different infection dynamics as immersed animals have additional physical layers of immune protection against the virus including the scales, skin, mucus layer, epithelial layer of the gills, and alimentary canal which all play fundamental roles in preventing pathogen entry into the body of the animal(Smith *et al*. 2019; Mokhtar *et al*. 2023). Another potential explanation is the difference in the crossing scheme, as Griot *et al*. (2021) used four full-sib backcross families (Eastern and Western Mediterranean sires, Western Mediterranean dams) and two farmed populations for their analyses. The relatively small number of families of diverse origin could explain the relatively low genetic variance explained by associated markers in the latter study.

As already mentioned, in all aforementioned studies, the precise identity of the putative causative variants remained elusive. Using imputed WGS data, Delpuech et al. (2023) identified four VNN-resistance QTLs on LG12 (presently Chromosome 3). The most significant one was centred on a region of approximately 50 kb (LG12:8,750,00-8,800,000), which includes two protein-coding genes (ZDHHC14 and IFI6/IFI27-like). Highly significant SNPs were represented by a non-synonymous substitution on ZDHHC14 and a non-coding polymorphism (the most significant one) 3.7 kb downstream of IFI6/IFI27-like. Additionally, all previous studies on the genetic assessment of VNN resistance in European sea bass(Palaiokostas *et al*. 2018; Griot *et al*. 2021; Vela-Avitúa *et al*. 2022; Delpuech *et al*. 2023) are based on a previous genome assembly (Tine *et al*. 2014), which was produced using short-read sequencing technologies, resulting in a highly fragmented assembly with several sequence gaps. Here, we used for the first time a novel, highly-contiguous genome assembly built using ultra-long sequence reads. In fact, analysis of the candidate QTL region in the old genome assembly used by Delpuech *et al*. (2023) shows that there are only two IFI6/IFI27-like genes in the candidate region, which likely correspond to LOC127358685 and ENSDLAG00005026832 in the novel assembly. Not surprisingly, in the old assembly there is a sequence gap between the ZDHHC14 locus and the two IFI6/IFI27-like genes in the region, the three missing IFI27-like genes that are part of the cluster in the new assembly are found in unplaced contigs in the previous version of the genome. In the novel assembly, the ZDHHC14 gene is located in chromosome 3, but at several Mb of distance from the region we identified as the most significantly associated with VNN-resistance. All this evidence suggests that the previous discordance regarding the position of the QTL could be explained, at least in part, by the lower quality of the previous genome assembly.

In our study, whole genome sequencing and imputation based on the novel genome assembly provided a clearer picture of the genetic background of VNN resistance in farmed sea bass when compared to the SNP array. The position of the major QTL was refined and narrowed to a relatively small genomic region, with the most significant SNPs overlapping two *IFI27L2A* genes, and one *IFI27L2* gene. The most significant SNP reported by Delpuech *et al*. (2023) is located 3.7 kb downstream of one of these genes (LOC127358685), putatively overlapping with the ENSDLAG00005026832 locus in the novel assembly. Co-localization of the most significant variants from GWAS and eQTL analysis strongly supports the hypothesis that the causative SNP(s) is(are) regulatory variant(s), significantly altering the expression of *IFI27L2* and the two *IFI27L2A* genes (Mancuso *et al*. 2019).

Human *IFI27L2A* encodes for interferon alpha-inducible protein 27-like protein 2A, which is an interferon-stimulated gene (ISG), and is sometimes referred to as ISG12, while *IFI27L2* which is also referred to as *ISG12B,* encodes for the interferon alpha-inducible protein 27-like protein 2. Upon detection of viral infection, host cells release interferons that in turn up-regulate the expression of several ISGs, which combat the infection through viral degradation and the inhibition of viral replication(Schoggins 2018). Interestingly, in mammals *IFI27L2A* has antiviral properties in the central nervous system (Lucas *et al*. 2016). Lucas *et al*. (2016) demonstrated, using a mice *IFI27L2A* knockout, that *IFI27L2A* protected mice against lethal West Nile virus (WNV) infection by significantly restricting replication of this virus in the different tissues of the central nervous system. Recently, a cluster of eight IFI27L2A homologues (ISG12.1-8) has been reported on zebrafish chromosome 13, and ISG12.1 was shown to suppress virus replication via targeting viral phosphoprotein(Guo *et al*. 2023), suggesting a conserved antiviral role of *IFI27L2A* in fish. *IFI27L2A* is up-regulated in response to VNN in European sea bass, and in fact an IFI27L2-encoding gene (FN665389.1), corresponding to LOC126358685, has been used since 2010 in several studies to monitor the response to either VNN infection or vaccination in this species(Nuñez-Ortiz *et al*. 2016). This is consistent with studies in other teleost species, where *IFI27L2A* was up-regulated in response to VNN infection (Toubanaki *et al*. 2022). The up-regulation of three (LOC127358630, LOC127358685, ENSDLAG00005026832) of the five interferon-induced genes present in the European sea bass in response to VNN, and the association of the VNN-resistance alleles with higher expression of these genes, strongly suggest that *IFI27L2A*/*IFI27L2* could be the causative genes underlying the major QTL for resistance to VNN. While the observed pattern appears to be tissue-and condition-specific for the three *IFI27L2A/IFI27L2* genes, this suggests that resistance is favoured by “constitutive” higher levels of mRNA of the genes.

The observed complex pattern of tissue- and condition-specific expression is not surprising, since duplicated loci are known to evolve through neo-functionalisation and sub-functionalisation (Lynch & Force 2000), leading to different tissue- and stimulus-specific patterns. Moreover, regulation of gene expression is a complex mechanism, involving the cooperation of multiple transcription factors and co-factor/co-repressors that recognise different DNA motifs (regulatory elements) located mostly in the proximity of the regulated gene, but that can also be located at a substancial distance of the gene promoter (Kim & Wysocka 2023). Finally, evidence from single-cell functional genomic studies has shown that “bulk” analysis on whole tissues and organ samples offers a coarse view of gene regulation (Carter & Zhao 2021). Albeit with a very limited sample size, single-cell midbrain transcriptomics in another VNN-susceptible species, the orange spotted grouper *Epinephelus coioides*, suggested that specific cell types are involved in the antiviral response and only certain neuronal cell lineages are damaged by the virus (Wang *et al*. 2021). While eQTL analyses using single-cell RNA-seq data are still prohibitively expensive for non-model species, they could vastly help us understand the observed expression patterns and genetic regulation of the different European sea bass IFI27L2/2A genes.

### Two SNPs in the promoter regions of IFI27L2A genes are strong candidate variants for the VNN resistance QTL

The combination of disease phenotypes, WGS genotypes, eQTL-level RNA sequencing, and ATAC-seq and ChIP-seq, significantly reduced the number of putative causative variants. In particular, the use of chromatin accessibility, an essential feature of regulatory elements, allowed us to narrow the number of candidates to two SNPs. Both are located on putative promoter regions of IFI27L2A genes, and are predicted to result in novel binding sites for two different transcription factors, KLF15 and TEAD4, which are involved in regulating gene expression in immune cells (El-Mayet *et al*. 2017; Luo *et al*. 2021). KLF15 is also an important stress-induced gene in fish (Carrizo *et al*. 2021), and experimental infection with VNN in European sea bass has been reported to activate a strong stress response (Lama *et al*. 2020). While evidence of the effect of genetic variants on TFBS predicted using human/mouse data should be taken with caution, high sequence conservation at the amino acid level frequently suggests that DNA-protein interactions are conserved. The two identified SNPs are strong causative candidates for the VNN-resistance QTL.

Population genomics data further supports the role of these two variants in resistance to VNN. It has been consistently shown that European sea bass individuals originating from Eastern Mediterranean populations have a significantly higher survival (up to three times) upon experimental infection with VNN compared to fish of Atlantic and Western Mediterranean origin [4, 7]. Our analysis of allele frequencies revealed that alternative alleles for both SNPs have a significantly higher frequency in Eastern Mediterranean populations compared to Atlantic and Western populations, further corroborating our findings. However, the difference is much more pronounced for the variants in CHR3:10077301, with a considerably higher frequency of the resistant allele, which suggests this variant is the strongest candidate causative mutation for the major QTL for resistance to VNN.

This CHR3:10077301 variant is located in the promoter of LOC127358685, generating the KLF15 binding site, and shows association with the expression of the three interferon-induced genes (LOC127358685, LOC127358630 and ENSDLAG00005026832) in both tissues. While it is located in the promoter region of LOC127358685, the variant could also modulate the expression of the other two genes, as promoters are known to often act also as enhancers (termed as Epromoters), exerting their effects on other loci(Andersson & Sandelin 2020; Malfait *et al*. 2023). However, we should not exclude the possibility that there could be other candidate variants, or even additional variants, are involved in the genetic basis of resistance; massive parallel reporter assays (MPRAs) have shown that at least 17.7% of eQTLs have multiple, tightly linked causal variants(Abell *et al*. 2022). Nevertheless, the SNP in CHR3:10077301 is a strong candidate to regulate resistance to VNN in European sea bass via modulation of the expression of the interferon-induced gene LOC127358685.

## CONCLUSIONS

Precise identification of causal variants for complex traits remains a major challenge in human and animal genomics, but integration of multiple functional evidence has already shown a great potential to finely map causative mutations. The integration of phenotyping, genome-wide genotyping and multiple functional genomics technologies assessing both the coding and non-coding regions of the genome have allowed us to identify the causative loci and mutation(s) underlying a major QTL for resistance to VNN in European sea bass. This study paves the way for biotechnological applications to generate fully VNN-resistant animals in European sea bass or other susceptible species; tools such as marker-assisted selection and genome editing that can exploit the discovered loci to increase VNN-resistance in aquaculture species, improving animal welfare and the sustainability of food production.

## METHODS

### Fish population generation and VNN challenge test

The experimental fish were produced in January 2020 using a commercial, NNV-free tested, broodstock from Valle Cà Zuliani Società Agricola srl (Pila di Porto Tolle, Rovigo, Italy). Dams were subjected to ovarian biopsy and those with a suitable stage of development of eggs were hormonally injected and stripped 72 h after the injection. Artificial fertilization was performed by mixing the eggs with previously stripped and preserved sperm(Fauvel *et al*. 2012). The mating scheme was based on a full-factorial design (25 dams × 25 sires). Eggs were mixed and distributed in four rearing tanks. At an age of 240 d post-hatching and an average weight of 13 grams, approximately 1,029 fish were transferred to the Istituto Zooprofilattico Sperimentale delle Venezie (IZSVe, Legnaro, Padova, Italy) and equally distributed into three close-system 2500-L tanks. Each tank was filled with artificial saltwater (30‰ salinity, temperature 21 ± 1°C, oxygen 6 ppm) and exposed to artificial photoperiod (10 h of light, 14 h of darkness). After an acclimation period of 14 days, fish were individually tagged with passive integrated transponder (PIT) tags and subjected to a VNN challenge test through intramuscular injection of 0.1 mL of a 1:100 dilution of the viral suspension (RGNNV 283.2009, stock titre 10^8.30^ TCID_50_ per mL(Panzarin *et al*. 2012). The procedure was performed on anesthetized animals (MS-222, 30 ppm). Water temperature was increased to 25 ± 1°C, while the other parameters (salinity, oxygen, photoperiod) were the same as in the acclimation period. Mortality was recorded twice a day. The challenge test ended after 29 d, when mortality returned to baseline levels. Resistance to VNN infection was defined as binary survival (BS, 0 survivor fish, 1 dead fish) and days to death (DD).

An additional group of fish (n = 324) from the same factorial crossing (i.e. full and half-sibs of the challenged animals) were exposed to VNN to assess the transcriptomic response to the infection. Of these 324 fish, 214 were VNN challenged through intraperitoneal injection of 0.1 mL of a 1:100 dilution of the viral suspension (RGNNV 283.2009, stock titre 10^8.30^ TCID_50_ per mL(Panzarin *et al*. 2012), while 110 were mock-challenged through injection of Dulbecco’s Modified Eagle Medium (DMEM), the cell culture medium where the virus was cultured. For each fish, brain and head kidney were sampled at 48 hours post-challenge for RNA sequencing (peak of the immune response, determined based on a preliminary transcriptomic study at 6h, 12h, 24h, 48h, 72h post-challenge on the same population; data not shown).

Fin clips of all the animals, including the parents, were collected and individually preserved in 70% ethanol and stored at 4°C for genotyping.

### DNA extraction and genotyping

The genomic DNA of all the fish used in this study (including 50 parents and 1353 offspring) was extracted from fin clips using Chelex method, employing Chelex as the extraction medium (Walsh *et al*. 1991) and genotyped on the bi-species MedFish SNP array (Peñaloza *et al*. 2021) containing 29,888 European sea bass SNPs, which was performed by IdentiGen Ltd (Dublin, Ireland). The array intensity files were imported to the Axiom analysis Suite v4.0.3.3 software for quality control (QC) assessment and genotype calling. SNPs were called following the default recommendations for diploid species (call rate threshold >97%). A total of 1,362 fish (97%) and 28,404 high-quality SNPs (95%) were retained for further analysis. The samples that passed QC included all 50 parental fish, 990 VNN resistance-phenotyped animals, and 322 VNN transcriptome challenge samples.

All parents (25 dams and 25 dams) and 40 resistance-phenotyped offspring were whole-genome sequenced by Novogene (Cambridge, UK) to obtain whole-genome genotypes for imputation. Genomic DNA was extracted from fin clip tissue using the salt-extraction method (Aljanabi & Martinez 1997). Subsequently, DNA sequencing libraries were prepared using Illumina TruSeq DNA Preparation Kits at the facilities of Novogene and then sequenced on a NovaSeq 6000 platform (Illumina, California, San Diego, USA) as 150PE reads to a sequencing depth of 15X. Raw sequence reads were assessed for sequencing quality using FastQC v0.11.9 (Andrews 2017), and consequently trimmed of any sequencing adaptors and low quality sequences using Trimmomatic v0.32 (Bolger *et al*. 2014); high quality reads of length ranging from 36bp to 150bp were retained for variant calling. Burrows-Wheeler Aligner (BWA) v0.7.187(Li & Durbin 2009) was then used to align the clean reads to the European sea bass genome dlabrax2021 (GCA_905237075.1) downloaded from Ensembl (http://ftp.ensembl.org/pub/release-107/fasta/dicentrarchus_labrax/dna/Dicentrarchus_labrax.dlabrax2021.dna.toplevel.fa.gz). Samtools v1.15 (Li 2011) and BCFtools(Li 2011) were subsequently used for genotype calling.

### Genotype quality control assessment and cleaning, parentage assignment and imputation

SNP array genotype data was further filtered using PLINK v1.9 (Chang *et al*. 2015), removing SNPs with genotype call rate across samples (geno) < 95%, minor allele frequency (MAF) < 5%, and significantly deviating from Hardy-Weinberg equilibrium (HWE) < 1E-4; samples missing > 10% of the genotypes were also removed from the analysis. After QC, 1,312 animals and 24,740 SNP markers were retained. Principal component analysis (--pca) was performed in PLINK v1.9 using all remaining SNPs to assess the genetic structure of the study population. Parentage assignment was performed using APIS package v1.0.1 (Griot *et al*. 2020) in R v4.1.1 through estimation of average Mendelian allele transmission probability between parents (sires and dams) and offspring using all 24,740 SNPs, allowing an assignment error rate of 1%.

WGS genotypes were filtered using vcftools v0.1.13 (Danecek *et al*. 2011), removing variants (SNPs and Indels) with more than two alleles (--max-alleles 2), genotyped in less than 50% of the samples (--max-missing 0.5), with genotype quality <30 (--minQ 30), low coverage <8 reads (--minDP 8) or very high coverage >31 reads (--maxDP 31.05), and with minor allele count <3 (--mac 3). Additionally, variants strongly deviating from Hardy-Weinberg equilibrium (--hwe 1E-7) in the parental population were also removed using PLINK v1.9. After QC, a total of 8,368,178 variants were retained for further analyses.

Fimpute version 3 (Sargolzaei *et al*. 2014) was used to perform genotype imputation for all the offspring (genotyped using the SNP array, 24,740 SNPs) using the parents as reference (genotyped using WGS, 8,368,178 SNPs), resulting in a dataset of ∼8.4M variants.

### Survival analysis and estimation of genetic parameters

Survival analyses were performed for the VNN resistance phenotyped offspring using the Survminer v0.4.9 package(Kassambara *et al*. 2021) in R v4.1.1. Bivariate restricted maximum likelihood (--reml-bivar) analyses were performed in GCTA version 1.93.2beta(Yang *et al*. 2011) to estimate genomic parameters (heritability and genetic correlations) of VNN resistance traits using SNP array and WGS imputed genotypes. VNN resistance phenotypes were preadjusted for population genetic substructure using the first and second principal components obtained using PLINK v1.9 as described above.

### Genome-wide association study

Genome-wide association study (GWAS) analyses were performed through single marker association testing using the mixed linear model leaving-one-chromosome-out (LOCO) implemented in GCTA v1.94.1 (--mlma-loco) (Yang *et al*. 2011). The following linear mixed model was fitted for the association analysis:

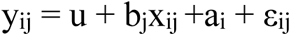

where y_ij_ was the pre-adjusted VNN resistance phenotypic value (binary survival or days to death) of the i^th^ individual, b_j_ is the allele substation effect of the j^th^ variant, x_ij_ are the j^th^ variant genotype classes coded as 0, 1, 2 (number of copies of the minor allele), a_i_ is the random additive polygenic effect (breeding value) of the i^th^ individual, and ε_ij_ is the random residual effect of the i^th^ individual with the x_ij_ genotype of j^th^ SNP. The random additive polygenic effects (a) are assumed to be independent and follow a normal distribution 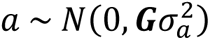 where 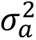 is the additive polygenic effect variance of the trait in the study population and *G* is the genomic relationship matrix between all individuals constructed in GCTA v1.94.1 (--make-grm) using the genotypes of all variants except those located on the chromosome of the variant under consideration. The random residual effects (ε) were also assumed to follow a normal distribution 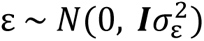 where 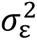 is the residual variance and ***I*** is an identity matrix of n x n dimensions (where n is the number of individuals in the analysis).

P-values for the variants from the analysis were adjusted for multiple testing using the Benjamini-Yekutieli (BY) method (Benjamini & Yekutieli 2001) and the variant was considered significantly associated with the trait at an adjusted P-value < 0.01. Phenotypic variance explained by a single variant was calculated as 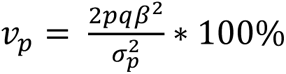, while additive genetic variance explained by each variant was calculated as 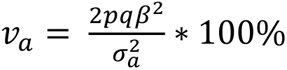, where p and q are the minor and major allelic frequencies of the variant, β is the allele substitution effect, 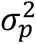 and 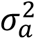 are the overall phenotypic variance and additive genetic variance of the trait in the studied population.

### RNA sequencing, differential gene expression and expression QTL analyses

Total RNA was extracted from the head kidney and brain of the 110 mock challenged fish and 214 VNN infected fish of the secondary experiment using RNeasy Mini Kit (Qiagen, Hilden, Germany) according to the instructions of the manufacturer. RNA concentration and quality was assessed using an Agilent 2100 Bioanalyzer with the RNA Nano 6000 kit (Agilent Technologies, Santa Clara, CA, USA), and only RNA samples with RIN >7 were processed for subsequent cDNA library construction and sequencing. cDNA libraries were constructed using the QuantSeq 3′ mRNA-Seq Library Prep Kit FWD (Illumina, Lexogen GmbH, Vienna, Austria) following manufacturer’s instructions, and sequenced on a NovaSeq 6000 sequencing platform (Illumina, California, San Diego, USA) as 75 bp single-end reads. Raw read sequence data was assessed for sequencing quality using FastQC v0.11.9 (Andrews 2017), and low quality reads and residual adaptors were removed using BBDuk v38.84 (Bushnell 2014). High-quality reads were then mapped against the European sea bass genome (GCA_905237075.1) using the short read aligner STAR v2.7.3a (Dobin *et al*. 2013). Gene counts were produced using featureCounts v2.0.0(Liao *et al*. 2014) and the annotation of the European sea bass genome, with the option “--primary” that counts the primary alignment of multimapping reads (in addition to uniquely mapped reads).

Differential expression between susceptible and resistant QTL genotypes was performed with the R package DESeq2 v1.38 (Love *et al*. 2014) using the standard pipeline. Briefly, separately for each tissue (i.e., brain and head kidney) and condition (i.e., mock and VNN-challenged animals), the relevant samples were extracted from the count table and imported in DESeq2 with the function ‘DESeqdataSetFromMatrix’; in order to remove uninformative genes that could contribute to background noise (Peruzza *et al*. 2018), the matrix of gene counts was filtered to keep genes with > 20 raw reads in total across all samples. Then, the filtered count table was normalised using the ‘varianceStabilizingTransformation’ function and differential expression analysis was performed using the function ‘DESeq’ by contrasting the resistant vs susceptible genotypes and performing a Wald significance test with adjusted p-value threshold < 0.05.

Additionally, using the VNN resistance phenotyped fish as reference, the genomic best linear unbiased prediction (GBLUP) approach implemented in GCTA v1.94.1 was used to compute genomic estimated breeding values (GEBVs) for resistance to VNN for the transcriptome immune challenge experiment (not phenotyped) using the linear mixed model bellow.*y* = *1_n_μ* + *Zg* + *e*

Where *y* is a vector of VNN resistance phenotypes (pre-corrected for genetic structure), ln is a vector of ones, *μ* is the overall average, *g* is a vector of random additive polygenic effects (genomic breeding values) assumed to follow a normal distribution 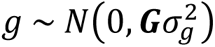 where G is the relationship matrix between all individuals constructed in GCTA v1.94.1 (--make-grm) using genome-wide markers, while 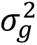 is the additive genetic variance of the trait, *Z* is a design matrix linking the random additive polygenic effects (genomic breeding values) in *g* to the phenotypes in *y*. And *e* is a vector of random residual values assumed to follow a normal distribution 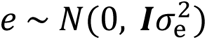 where ***I*** is an identity matrix and 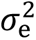 is the residual variance. Pearson’s correlations were then computed between gene expression log_2_CPM values of expressed genes and VNN resistance GEBVs for mock challenged and VNN infected groups and each tissue separately using Hmisc version 4.7-0 package in R v4.1.1 (Harrell Jr & Harrell Jr 2019). Subsequently, based on the differential gene expression and correlation analysis results, GCTA v1.94.1 LOCO (--mlma-loco) was used to identify expression QTL for the candidate VNN-resistance genes in each tissue and condition separately.

### Functional characterization of the VNN resistance QTL region in the European sea bass genome

Multiple functional genomics datasets (including RNA-seq, ATAC-seq and ChIP-seq) generated from naïve juvenile seabass treated with Polyinosinic:polycytidylic acid [poly(I:C)], a synthetic immunostimulant used to simulate double stranded-RNA (dsRNA) viral infections, were used to functionally annotate the VNN resistance QTL region in the European sea bass genome. Briefly, individuals were stimulated by intraperitoneal injection with PBS or Poly(I:C) (6 replicates/condition). Animals were sacrificed 24h post-infection (hpi) and head kidney sampled for RNA-seq, ATAC-seq and ChIP-seq library preparation (6 replicates/condition). RNA-seq, ATAC-seq and ChIP-seq libraries were also produced for brain of healthy juvenile sea bass (3 males and 3 females). All these procedures were part of a large annotation effort under the framework of the EU project AQUA-FAANG (https://www.aqua-faang.eu/).

For all the above-mentioned samples, total RNA was extracted using RNeasy Mini Kit (Qiagen, Hilden, Germany) according to the instructions of the manufacturer, and high quality RNA was then sent to Novogene for cDNA library preparation and sequencing on the Illumina NovaSeq 6000 system platform (Illumina, California, San Diego, USA) as 150 bp paired-end reads.

From the same samples, ATAC-seq libraries were constructed using the Omni-ATAC protocol (Corces *et al*. 2017) with species and tissue-specific modifications (details here: https://data.faang.org/api/fire_api/experiments/NMBU_SOP_OmniATAC_protocol_20200429.pdf). Briefly, frozen tissue fragments (20-30 mg) were homogenized by using a pre-chilled 2-ml dounce tissue homogenizer containing homogenization buffer. Connective tissue and residual debris were pre-cleaned by filtration through a 70-μm nylon filter. Intact nuclei were then collected by density gradient centrifugation over iodixanol (25% iodixanol, 29% iodixanol, 35% iodixanol). For brain samples, after PBS washing, nuclei were counted by trypan blue staining and 50,000 nuclei were then recruited for Tn5 transposase reaction by using Illumina Tagment DNA Enzyme and Buffer kit (Illumina, California, San Diego, USA). For head kidney samples, all washing steps were skipped in order to avoid nuclei aggregation and 1 µl of nuclei band was directly employed for transposase reaction. Transposition reactions were cleaned up with MinElute PCR Purification Kit (Qiagen, Hilden, Germany) and uniquely barcoded libraries were obtained following amplification using Illumina Nextera DNA Unique Dual Indexes (Illumina, California, San Diego, USA). After quality assessment, all libraries were sent to Novogene (Cambridge, UK) for 150bp paired-end sequencing on the Illumina NovaSeq 6000 system platform. Sequence data analyses were performed using the NF-core pipeline for ATAC-seq analysis (nf-core/atacseq v1.2.1, https://github.com/nf-core/atacseq).

Following the manufacturer’s instructions, ChIP-seq libraries for the same samples were prepared using the µChIPmentation Kit for Histones (Diagenode). Three histone marks were investigated: H3K4me3 (promoter regions), H3K27ac (active enhancer and promoter regions) and H3K27me3 (associated with Polycomb repression complexes). After quality assessment, all libraries were sent to Novogene and sequenced by Illumina NovaSeq6000 150PE in order to obtain 45M reads per sample. All sequencing data analyses were conducted through the NF-core ChIP-seq analysis pipeline (https://github.com/nf-core/chipseq).

Additionally, a total of 20 fish (10 mock- and 10 VNN-infected) from the transcriptome immune challenge experiment were used to assess chromatin accessibility in the brain and head kidney tissue under mock and VNN infection conditions. Forty ATAC-seq libraries were constructed from frozen tissue using the procedures described above, and sequenced by Novogene (Cambridge, UK) as 150 bp paired-end reads using the Illumina NovaSeq 6000 platform. This data was also analysed using the NF-core pipeline for ATAC-seq analysis (nf-core/atacseq v1.2.1, https://github.com/nf-core/atacseq).

For each sample and condition (i.e., PBS-stimulated HKs, Poly(I:C)-stimulated HKs, brains from male and female individuals), ATAC-seq and ChIP-seq data were combined to infer genome-wide chromatin state and associated regulatory elements using ChromHMM v1.24 (Ernst & Kellis 2017), assuming 10 different chromatin states. This information was crossed with genetic variation in the VNN resistance QTL region, and the putative impact of variants overlapping regulatory elements on putative transcription factor binding sites was predicted using FABIAN-variant (Steinhaus *et al*. 2022). Briefly, 41 bp sequence fragments centred around each variant were analysed on the web-based version of the software (v2022-05-26), with the transcription factor flexible models (TFFMs)-detailed model option.

### Comparative genomic sequence analysis

The Interferon Alpha Inducible Protein 27-like genes (*IFI27L2A* and *IFI27L2*) encoding protein sequences in the European sea bass genome (GCF_905237075.1), the striped bass (*Morone saxatilis*) genome (GCF_004916995.1), and the gilthead sea bream (*Sparus aurata*) genome (CA_900880675.1) were aligned using MUSCLE (Edgar 2004). Striped bass is phylogenetically very close to European sea bass (same family, Moronidae), while gilthead sea bream is phylogenetically more distant. From European sea bass, sequence XM_051391909.1 (encoded by locus LOC127358638) was too divergent to be reliably aligned with the other IFI27L2/2A transcripts and hence was removed from the analysis. Multiple alignment of the remaining sequences was cleaned using Noisy v1.5.12 (Dress *et al*. 2008)[70]. The final alignment was used as input for reconstructing a gene tree using maximum likelihood (PhyML+SMS) and Bayesian methods (MrBayes). To assess the genomic evolution conservation of the *IFI27L2/2A* genes between European sea bass and striped bass multiple sequence alignment of the genomic regions harboring these genes in two species (European sea bass: Chromosome 3: 10,075168 – 10,097,914; Striped sea bass: Chromosome X: 15,596,712 – 15,606,764) was performed using mVISTA (Frazer *et al*. 2004). Additionally, to assess the gene evolutionary similarity between the five copies of the IFI27L2/2A genes in European sea bass multiple sequence alignment of the protein sequences of the five IFI27L2/2A gene loci in the VNN QTL region was performed using MEGA11 v0.10 software (Tamura *et al*. 2021) and visualized using a Bioconductor R package ggmsa v1.3.4 (Zhou *et al*. 2022)

### Availability of data and materials

Whole genome imputed genotypes and phenotypes have been submitted to the Europeans Variant Archive (EVA) under the accessions PRJEB77213 and ERZ24798734. Whole-genome raw sequencing data have been uploaded to the NCBI Short Read Archive (SRA) under BioProject PRJNA1110973. RNA-sequencing data have also been uploaded to the NCBI SRA under BioProject accession number PRJNA1122486. ATAC-seq and ChIP-seq sequencing data, and functional annotation from head kidney samples have been uploaded to the EMBL-EBI repository under accession numbers PRJEB52284 and PRJEB59557, respectively. ATAC-seq and ChIP-seq sequencing data from brain samples have been uploaded to the EMBL-EBI repository under accession numbers PRJEB52775 and PRJEB59432, respectively.

## Acknowledgements

We thank Valle Cà Zuliani Società Agricola srl for providing the fish used for the VNN challenge experiment.

## Author Information

Conceptualization, LB, DR and RDH; disease challenge, DB, AF,FP; data generation, SF, CP,AT, FP,RF; analysis and visualization, RM, SF, LP, GDR, MB,SF, CP, LB, DR; funding acquisition, LB, RDH and CT; project administration, RDH, LB and DR; supervision, CT, RH, LB, DR; writing, RM, LB and DR. All authors edited and reviewed the manuscript.

## Ethics declarations

### Ethics approval and consente to participate

Animal care and was fully compliant with the prescriptions of the Directive 2010/63/EU of the European Parliament and of the Council, implemented at national level through the D. Lgs 4 March 2014, n.26. All experiments and VNN challenge protocols were authorized by the Italian Ministry of Health (auth. 975/2016-PR).

### Consent or publication

Not applicable.

